# Comprehensive contact tracing during an outbreak of alpha-variant SARS-CoV-2 in a rural community reveals less viral genomic diversity and higher household secondary attack rates than expected

**DOI:** 10.1101/2023.11.17.567570

**Authors:** Audun Sivertsen, Nicolay Mortensen, Unni Solem, Eivind Valen, Marie Francoise Bullita, Knut-Arne Wensaas, Sverre Litleskare, Guri Rørtveit, Harleen M. S. Grewal, Elling Ulvestad

## Abstract

Sequencing of SARS-CoV-2 genomes throughout the COVID-19 pandemic has generated a wealth of data on viral evolution across populations, but only a few studies have so far explored SARS-CoV-2 evolution across transmission networks of tens to hundreds of persons. Here, we couple data from SARS-CoV-2 sequencing with contact tracing data from an outbreak with a single origin in a rural Norwegian community where samples from all exposed persons were collected prospectively. A total of 134 nasopharyngeal samples were positive by PCR. Among the 121 retrievable genomes, 81 were identical to the genome of the introductor, thus demonstrating that genomics offers limited additional value to manual contact-tracing. In the cases where mutations were discovered, five small genetic clusters were identified. We observed a household secondary attack rate of 67%, with 92% of household members infected among households with secondary transmission, suggesting that SARS-CoV-2 introduction into large families are likely to affect all household members.

**Importance:** In outbreak investigations, obtaining a full overview of infected individuals within a population is seldom acheived. We here present an example of just that, where a single introduction of B1.1.7 SARS-CoV-2 within a rural community allowed for tracing of the virus, from an introductor via dissemination through larger gatherings, into households. The outbreak occurred before widespread vaccination, allowing for a “natural” outbreak development with community lock-down. We show through sequencing that the virus can infect up to five consecutive persons without gaining mutations, thereby showing that contact tracing seems more important than sequencing for local outbreak investigations. We also show how families with small children are less likely to contain spread to all family members if SARS-CoV-2 enters the household either by a child or a caregiver, as isolation of the primary infected is difficult in such scenarios.

## Introduction

Coronavirus infectious disease-2019 (COVID-19) is caused by the severe acute respiratory syndrome coronavirus 2 (SARS-CoV-2). The disease emerged in December 2019 in Wuhan, China, and was declared a pandemic by the World Health Organization (WHO) on March 11^th^ 2020. Several strategies, including manual tracing of contacts of patients with positive PCR-results and investigations of viral genome variability in samples from patients infected with SARS-CoV-2, have been advocated to curb the spread and to track the routes of transmission during outbreaks (1–3).

Although important for restricting viral spread, the strategies are prone to miss asymptomatic patients who often remain untested and patients who present with symptoms before viral shedding has reached the threshold for PCR-detection (4). Furthermore, elucidation of the transmission chain has been hampered by the low mutation rate of SARS-CoV-2, the high frequency of identical genomes across several links of transmission, and variations in methods of sampling. While retrospective studies identifying symptomatic cases report household secondary attack rates (HSAR) frequencies of 25-50% (5), prospective sampling with symptomatic and asymptomatic cases exceed 90% (4), especially in households with many residents (6). Transmission mapping limitations may be overcome by coupling data from viral genome variability to epidemiological investigations (7–10) and through modelling supported by large sets of metadata (11–13).

We here present a cluster of 134 persons infected with SARS-CoV-2 from a single origin in a relatively confined population. Using data from highly curated manual contact tracing we compare transmission routes with occurrence of mutations in the outbreak strain and also estimate the HSAR of the affected households.

## Results

### Population characteristics, SARS-CoV-2 testing and contact tracing

Ulvik is a municipality in Western Norway with 1080 inhabitants, one school, one kindergarten, and one nursing home. On January 25^th^ 2021, Ulvik experienced an outbreak of SARS-CoV-2. Prior to the outbreak, only seven cases of SARS-CoV-2 had been detected and only institutionalized elderly patients had been vaccinated against COVID-19. The patterns of housing, congregations, and close encounters in rural Ulvik are well-defined, thus facilitating contact tracing. Nasopharyngeal sampling, transportation of samples to Haukeland University Hospital, PCR testing and test result deliverance to the health authorities in Ulvik was done within 1-2 days with most samples.

A total of 809 samples for testing were obtained from 554 unique persons during the outbreak period, of which 134 persons were confirmed as SARS-CoV-2 positive. Twelve percent of the population was infected, a large proportion of whom were children and young adults. The chain of contagion was identified by contact tracing. The Chief county physician and other health personnel kept track of who likely infected whom progressively as each infectee tested positive for SARS-CoV-2. Contact tracing also included data on the most probable arena of infection, who lived with whom in households, as well as the age of the infected. A selection of metadata is present in supplementary table 1, and transmission links are present in supplementary table 2.

### SARS-CoV-2-transmissions in school and kindergarten – adults introduce, children disseminate

Dates of sampling and reports of positive tests to the health authorities are depicted in Figure 1A. The source patient contracted COVID-19 in Italy but did not display symptoms until after the tenth day of the quarantine period. The SARS-CoV-2-positive test reports and contact tracing obtained on day one of the outbreak, a Friday, made evident that the virus had been transmitted to both the kindergarten and the school by household members of the source patient. All individuals in the kindergarten and in two school classes were quarantined. At day 4 of the outbreak (Monday), contact tracing suggested that 7 out of 10 school classes had been exposed to SARS-CoV-2, and the local health authorities decided to close the school on day four of the outbreak. The same day, the laboratory confirmed that the transmitting SARS-CoV-2 virus was an alpha variant (B1.1.7).

**Figure 1.**
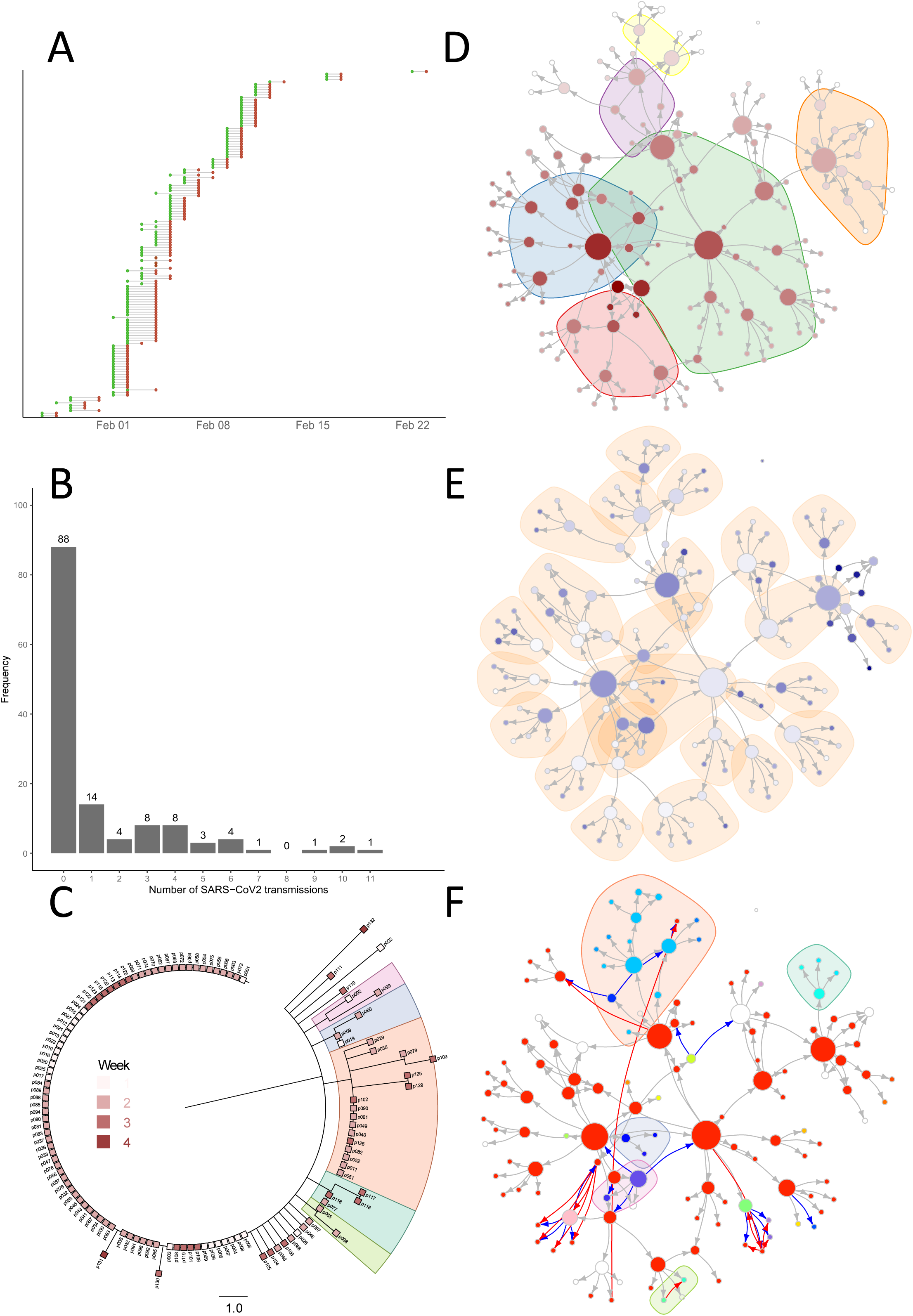
depicts the contact tracing network with associated metadata. Each node represents a person, and node size corresponds to the likely number of infectees (represented as directed edges) originating from the node. A: Lollipop chart showing dates of sampling (green dots) and feedback of a positive test result (red dot) for all persons in the dataset. B: Bar chart quantifying how many patients that infect a given number of others, by extracting the number of outbound edges from all nodes in the contact tracing network. C: Unrooted phylogenetic tree of all 121 available SARS-CoV-2 genomes as squares which are colored by sampling date from the 1st week (pale red) to the 4th week (dark red) of the outbreak. Underlying shades highlight genetic clusters corresponding to underlying shades in figure 1F. Scale shows number of mutations from source. D-F: Stylised transmission networks. Nodes represent persons within the network, and node size refers to the number of connected edges/transmissions. D: Transmission across zones. The nodes are colored by place in the transmission chain, where the source node is shaded the darkest red and the last infectees are pale red. Transmission clusters outside households (two kindergarten zones, two school classes, a youth club, and an elderly care center) are highlighted in colored shades underneath the network. E: Transmission within households. Nodes are coloured by shades of blue where older patients have a deeper shade of blue as compared to younger patients. Underlying shades show households. F: Variant calling of virus genomes resulted in 28 distinct genotypes with one or more SNPs. Each node is colored with divergent colors representing each distinct genotype. The source strain appears bright red. Phylogenetic clusters from common informative SNPs appear as colored shades underneath the network. Edges colored blue represent transmissions made likely by manual tracing but unlikely given genomic data. Red edges are novel transmission links identified through retrospective interrogations of contacts between patients given genomic data.

In this outbreak, four persons transmitted SARS-CoV-2 to nine or more persons, thereby acting as “superspreaders” in our network (Fig. 1B, also corresponding to the four largest nodes in Fig. 1D-F). Three of these introduced SARS-CoV-2 in the kindergarten and school. One “superspreader” also worked at the nursing home, thereby causing an outbreak among residents and employees. Transmission of SARS-CoV-2 was also detected at the local youth club. Transmission also occurred within households, where infectors had access to fewer infectees (Fig. 1B).

Fig. 1D shows a stylized version of the transmission network derived from the comprehensive tracing of contacts, with transmission arenas outside households as colored shades underneath the network. Each node represents a case, and the size of the nodes reflects the number of connected edges. Edge direction shows likely transmission events based on contact tracing. Some persons may have several potential infectors.

By adding metadata to the transmission network, we were able to identify the likely transmission routes within families as depicted in Fig. 1E, where underlying shades correspond to households. All nodes are colored in shades of blue by age with the oldest in the deepest shade. The age distribution of cases ranged from 2 to 99 years.

### SARS-CoV-2 sequences do not suffice to specify transmission events

After filtering out sequences with poor coverage, 121 SARS-CoV-2 genomes contained sufficient genetic information to be included in a phylogeny analysis by Nextstrain. Eighty-one (67%) of the 121 viral genomes were identical to that of the source patient (hereafter called source strain to distinguish it from other isolates with mutations). They could, therefore, not be used to validate transmission events, see unrooted phylogeny in Fig. 1C and color of nodes in Fig. 1F. As depicted in Fig. 1F, the virus could pass through up to five transmissions without displaying mutations. Of the remaining genomes, some had shared SNPs, thereby enabling the identification of five clusters; see correspondingly colored underlying shades in the phylogenetic tree (fig. 1C) and in the transmission network depicted in Fig. 1F.

A comparison of the manually curated transmission network and the network based on genomic information revealed some contradictory transmission routes. In all cases but one where the transmission arrow was transposed by addition of genomic data, the change originated from a mutated genome being present within a transmission chain in which infectees carried the source strain. By performing local re-interrogations of contact points in these cases, some alternative transmission routes could be added (red edges, Fig. 1F), and others could be subtracted (blue edges, Fig. 1F).

Two families had a primary infector whose SARS-CoV-2 genome contained a mutation that did not propagate to other household members of which several had the source strain, thus necessitating the identification of alternative household index patients.

In another case, a child did not receive the SARS-CoV-2 virus through household transmission of a mutated variant, but instead received the source strain from attendance in kindergarten. In a third case, a previously undisclosed longer conversation led to transmission of the source strain, where the earlier presumed transmitter carried a mutated strain which was unlikely to revert back to the source strain in the infectee.

In a fourth case, a cluster arose within a single family, where the infector had the source strain and three of four infectees carried strains with mutations. Two of the three mutated strains shared a common SNP. It is unclear whether the two connected mutated strains represented independent transmission between these two family members, or if the SARS-CoV-2 strain of the household index patient had hypermutability throughout the infective period.

The only unresolved discrepancy between epidemiological and genomic data was a single strain (Fig. 1F, blue node with inbound blue edge at the lower right side) where genomic data pointed to inclusion within the top cluster (orange shade, Fig. 1F), but no epidemiological connection could be found. This particular SARS-CoV-2 genome suffered from relatively poor coverage (median 652) and alignment to the reference strain (82,8%) and we can not exclude base calling errors as a confounding factor.

### Dense sampling gives a household secondary attack rate of 77%

In addition to the 130 infectees which have a known family constellation (4 institutionalized patients or single dwellers are not included) (Fig. 1E), 25 additional household members did not test positive for SARS-CoV-2 despite performing repeated nasopharyngeal swabs for PCR analysis. These persons are not shown in the network.

Thirty-eight households in the network consist of at least two persons, and comprise 155 residents In total. Of these households, 74% (28/38) experienced secondary transmissions. Assuming a single household index case per household (38 in total), the secondary attack rate in this population is 77% (92/117). Limiting the denominator to include only members of households with secondary transmission, the secondary transmission rate is 92% (92/100). This indicates the extent of secondary transmission if a household was not able to contain its index case. The feasibility of containment measures is further demonstrated by examining the 13 households where the household index case is a child. Only 1/13 of these households (8%) were able to limit further spread of the virus, as opposed to 9/25 (36%) households with an adolescent (≥13 years) or older as the household index case (p=.060). Figure 2 illustrates how households with secondary transmission more often have household index cases in the lower end of the age spectrum. The size of the dots reflects the number of people in each household. Those households where secondary transmission did not take place were characterized by being smaller (2-3 persons) and consisted of adults living with other adults and/or with teenagers.

**Figure 2.**
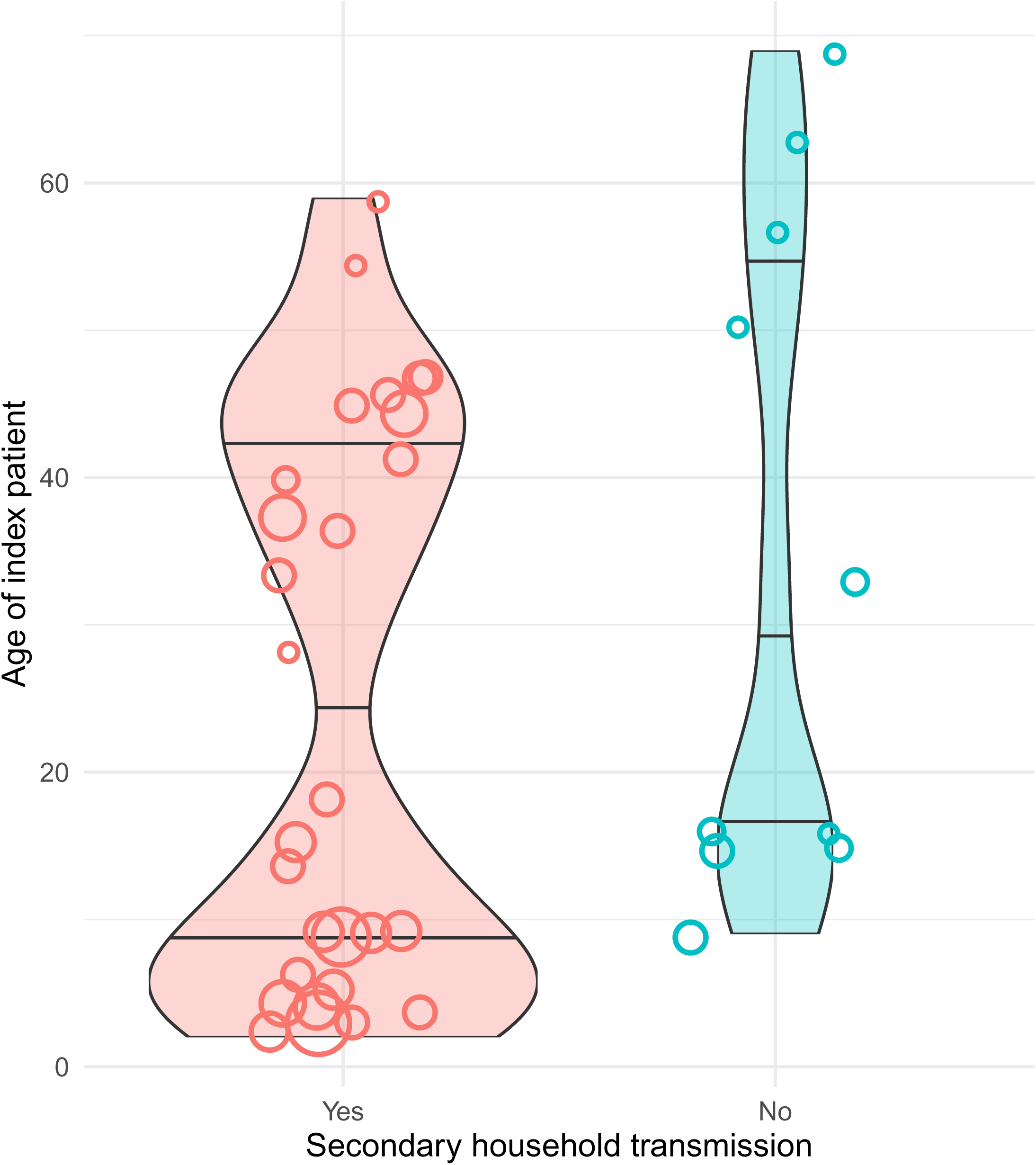
shows the household index patient’s age (y-axis) in households with and without secondary transmission (x-axis). Each circle represents a household, and the width represents household size. Violin plots show the age distribution, in which violin width is normalized to count and reflects the number of households with household index patients in the given age range.

## Discussion

We present the dynamics of a large connected outbreak of COVID-19 caused by the SARS-CoV-2 B.1.1.7 strain. The outbreak occurred in a local community, thus demarcating it from previously reported densely sampled outbreaks occurring in healthcare institutions (14, 15). The ecological settings and transmission dynamics are therefore distinct, especially since contact between households is likely less prevalent than contact between rooms in a ward or between wards in a health institution. The outbreak occurred before population-wide vaccination took place, thus providing a glimpse of SARS-CoV-2 propagation throughout an immunologically naïve population.

Of the 134 PCR-positive samples, 121 sequences could be extracted to create consensus sequences. Of these, 81 sequences were identical, and these identical sequences could be retrieved through as many as five transmission links across six individuals. This underscores that genomics does not provide enough resolution to resolve SARS-CoV-2 transmission on such a granular scale as displayed here or by others (8, 10, 13, 16). We chose to abstain from calculating minor variants as they seem to exist between a minority of transmission pair SARS-CoV-2 genomes, have limited ability to inform transmission directionality and apparently most often arise *de novo* within infected patients (17–19). Given the mutation frequency of SARS-CoV-2 compared to other viral pathogens, which is not exceptionally high (20), we presume that viral genomics in densely sampled outbreaks, in general, may show similar limited utility with transmission chain resolution compared to epidemiological data. Our sequencing results are consistent with results obtained during a large community outbreak of the Delta (B.1.617) variant in China (n=167) (10). A similar absence of genetic variance between samples was found, with most SARS-CoV-2 sequences being identical with occasional clusters of isolates with singular or common SNPs. However, the study did not report the vaccination status of infected persons nor the HSAR, thus limiting the results for resolving the transmission dynamics in naïve populations.

We observe a high HSAR compared to other estimates (5). There are several reasons why we believe we were able to obtain accurate estimates of HSAR in this outbreak. First, the calculation of HSAR relies on identifying all cases in an exposed population, regardless of symptom presence or varying incubation time across infected individuals. As such, infrequent, symptom-based or delayed sampling could limit case detection when utilizing PCR. Retrospective case identification by serology fails to identify all cases (21). Second, In contrast to others (22), we did not exclude households where several persons may have introduced SARS-CoV-2 into the household, as this occurred in a minority of cases (e.g. twins who attended the same class in school). As this community is interconnected, contagion could emanate from several sources, as corrected transmission pathways in Fig. 1F show. As these hidden “cross transmissions” are likely to occur also in other datasets and are hard to correct without accompanying metadata, we still believe that HSAR rates calculated here are representative.

Multiple circulating lineages introduced into the community at once may also disrupt contact tracing and assumptions of transmission directionality. Iterative progress in study design of HSAR studies during the continuation of the pandemic may have resulted in better capture of infected individuals, thus downplaying secondary attack rate in early variants. In population surveillance settings, identifying the denominator would be difficult, as identifying the number of exposed persons is uncertain for many COVID-19 cases.

Although children were the drivers of transmission in this outbreak by accounting for a larger proportion of people infected than adults, three out of the four “superspreaders” who transmitted SARS-CoV-2 to many others were adults, and likely reflects that adults more often than children move between different social arenas in the network.

We observed several instances of successful household isolation where no other household member got infected, particularly in households with few persons and adolescent or adult household index persons. In contrast, households with many members and children are not able to contain transmission within the household. As the majority of the drivers of infection within households are children or caregivers of children, our data shows that children are more likely to transmit SARS-CoV-2 within households compared to adults. A meta-analysis comparing SARS-CoV-2 transmission from adults and children found similar secondary transmission rates of SARS-CoV-2 among the groups (23). Our findings may then be explained by the demography of the population, with many large households with small children who can not be physically separated from their families.Households with secondary transmission often have many members, which includes children (as shown in Fig 2). Therefore, the exclusion of households with potentially more than one primary infector will bias results towards lower HSAR.

In conclusion, the data gathered by this single outbreak shows that genomic sequencing of SARS-CoV-2 is not likely to inform transmission events in connected clusters, but may occasionally provide extra data when mutations occur. During the outbreak, HSAR rates are affected by how many people are living within the household. If children introduce SARS-CoV-2 into a household, transmission often involves all other household members.

Some limitations exist - the capture of asymptomatic cases is still an issue, and during the outbreak, no standardized research protocol for sampling exposed individuals was in place. Variability in testing compliance, especially when testing at multiple timepoints, is to be expected. Albeit - testing all exposed persons at least once at a time point where the likelihood of obtaining a positive test is high, in addition to good compliance for testing in persons with either symptoms or possible exposure, suggests that most infected cases were discovered. Hidden transmission would likely cause symptomatic disease locally outside known paths of exposure - only one positive test originated from an asymptomatic person without a known link to any infector.

## Material & Methods

### SARS-CoV-2 RNA-extraction and PCR

Nucleic acids from the nasopharyngeal sample were extracted on the MagnaPure96 platform (Roche, Mannheim, Germany) using the MagNA Pure 96 DNA and Viral NA Large Volume Kit (Roche). The in-house real-time PCR was based on the E-gene primers and probes as described by Corman et al. (24) and was run on a LightCycler 480 instrument (Roche) in a 20 µl reaction volume using the QuantiNova Pathogen mastermix (Qiagen, Hilden, Germany). Ct values for positive samples are given in supplementary table 1.

### SARS-CoV-2 RNA-sequencing

Viral RNA from 127 of 134 total samples were sequenced by the Illumina NovaSeq6000 (Illumina, San Diego) with 150 bp paired-end reads. The library was generated by Swift Normalase Amplicon Panel (SNAP) SARS-CoV-2 with Additional Genome Coverage. The pipeline used for sequence quality assessment, variant calling and consensus sequence generation is available at https://github.com/nsc-norway/COVID-19-seq. Four genomes were sequenced at the Norwegian Institute of Public Health using the Oxford Nanopore GridION (Oxford Nanopore Technologies, Oxford), with preceding sample preparation using the ARTIC protocol v.2 (25). Variant calling and consensus sequence generation of Illumina sequences were done with iVar (26). The phylogeny from nextstrain.org (27) was used to define clusters. Three samples were lost during sample retrieval. All sequence reads are available in ENA BioProject PRJEB65109, with individual sample accessions provided in supplementary table 1.

### Household secondary attack rates (HSAR)

The HSAR was estimated by dividing the number of infected household members, (not counting the person introducing the infection to the household) by the total number of household members eligible to be infected by the household index person. Pearsońs χ^2^ was used for comparison of HSAR in households with secondary transmission and a child as the primary infector as opposed to an adult. A child in this setting was defined as a person below 13 years.

### Computational analyses

Figures and network plots were made in R, using igraph, GGally, RColorBrewer and ggplot2 packages (28–30).

### Ethics

The study was approved by the Regional Committee for Medical and Health Research Ethics in Western Norway (REK ID: 248964) and performed in accordance with the Declaration of Helsinki. Written informed consent was obtained from all participants or their legal guardian/close relative at the time of recruitment. Specific information regarding the introduction of SARS-CoV-2 into the community and outbreak containment through closures of school and kindergarten are publicly known through mass media communications.

## Acknowledgements

We thank the team of health personnel at the Ulvik municipality for assisting with contact tracing and greatly improving data quality. We also thank the Norwegian Public Health Institute for sequencing four SARS-CoV-2 strains and sharing sequence data.

## Supporting information

**Table S1: Metadata of patients, PCR and sequencing.** Contains pseudo-ID of patients. Ct values of SARS-CoV-2 PCR (Ct), if Ct is 0, the PCR was performed on a platform which does not report Ct values. Genotype (28 total genotypes). Cluster refers to genetic cluster of SARS-CoV-2 genomes. Platform refers to which sequencing platform the SARS-CoV-2 genomes were sequenced on – Illumina NextSeq, Oxford Nanopore GridIon, or not sequenced. Qual refers to iVar assessment as an aggregate of genome coverage and depth. WGS_mean and WGS_median sequencing depth. ACCESSION is sample ID in ENA, ENA_ALIAS is a searchable term in the ENA database. Information of household membership and exposure through specific infection arenas is available upon request.

**Table S2: Contact tracing data**. Contains edge information used to create network seen in Fig. 1D-F. Accompanying colors dark grey (edges uncorrected by SARS-CoV2 sequencing), blue (edges subtracted by SARS-CoV2 sequencing) and red (edges added by SARS-CoV2 sequencing). Network topology was created with edges without the color “red”.

